# Exploring 3-Aminobenzoic Acid as a Therapeutic Dietary Component for Enhancing Intestinal Barrier Integrity in Ulcerative Colitis

**DOI:** 10.1101/2024.08.18.608525

**Authors:** Miho Tanaka, Takahiko Toyonaga, Fumiyuki Nakagawa, Takeo Iwamoto, Akira Komatsu, Natsuki Sumiyoshi, Naoki Shibuya, Ayaka Minemura, Tadashi Ariyoshi, Asami Matsumoto, Kentaro Oka, Masayuki Shimoda, Masayuki Saruta

## Abstract

**Background:** Dietary components and their metabolites produced by intestinal bacteria play a crucial role in maintaining intestinal epithelial integrity. Disrupted epithelial integrity increases permeability and leads to chronic inflammation in the colon, known as ulcerative colitis (UC), in genetically predisposed individuals. However, the gut microbial metabolites regulating epithelial permeability remain unexplored and their metabolism in UC patients is unclear.

**Methods:** A library of 119 gut microbial metabolites was screened for their ability to reduce epithelial permeability in Caco2 cell monolayers. The diet containing 3-aminobenzoic acid (3-ABA) was identified using liquid chromatography with quadrupole time-of-flight mass spectrometry. The abundance of fecal 3-ABA was compared between UC patients and healthy individuals followed by 16S rRNA metagenomic analysis to estimate the gut microbial function in ABA degradation. The anti-inflammatory effect of 3-ABA was examined in a mouse model of dextran sodium sulfate-induced colitis.

**Results:** Stimulation with 3-ABA reduced epithelial permeability and enhanced barrier integrity in Caco2 cells by modulating the tight junctional regulatory pathway. 3-ABA was abundant in beans and decreased in the feces of UC patients. Functional prediction analysis of gut microbiota revealed an accelerated degradation of ABA with significant up-regulation of *mabA*, a gene encoding a bacterial enzyme involved in 3-ABA degradation, in UC patients. Rectal and oral administration of 3-ABA ameliorated experimental colitis in mice.

**Conclusion:** 3-ABA abundant in beans enhanced intestinal epithelial integrity and ameliorated experimental colitis in mice. Proactive intake of 3-ABA might be a novel treatment approach for UC.

## Introduction

The intestinal tract is a unique organ where the host coexists harmoniously with microorganisms. This delicate host-microbe symbiosis relies on a single layer of intestinal epithelial cells (IECs) which serves as a physical barrier separating the mucosal immune system from harmful bacteria through tight junctional connections. The epithelial barrier breakdown compromises this symbiosis, allows the invasion of luminal bacteria, and results in intestinal inflammation. Therefore, the integrity of intestinal epithelium is tightly regulated for maintaining gut health. Recent studies have highlighted the crucial role of dietary components and their metabolites produced by intestinal bacteria in regulating intestinal epithelial integrity[1, 2]. Notably, the Western pattern diet characterized by high intakes of refined grains, red meat, and high-fat dietary products has been shown to compromise intestinal epithelial integrity[3]. Dysbiosis, defined as compositional changes and decreased diversity in gut microbiota, further disrupts intestinal epithelial homeostasis by altering microbial metabolites[1]. Such detrimental changes increase IEC permeability and raise the risk of ulcerative colitis (UC), a chronic inflammatory condition of the colon, in genetically predisposed patients^3^.

UC is one of the two major forms of inflammatory bowel disease (IBD) along with Crohn’s disease and is characterized by relapsing and remitting symptoms, such as increased stool frequency, rectal bleeding, and abdominal pain, necessitating lifelong medical treatment[4]. While increased IEC permeability is a key trigger of colonic inflammation in UC, current therapies are primarily designed to target immune cells or immune-related pathways with limited focus on directly enhancing intestinal epithelial integrity. An exception is 5-aminosalicylic acid (5-ASA), the first-line drug for UC, which is believed to exhibit an anti-inflammatory effect via activation of peroxisome proliferator-activated receptor (PPAR) in IECs[5]. With many patients showing inadequate responses to current therapies and facing increased risks of infections or cancers due to immunosuppressive treatments, there is an unmet need for novel therapeutic strategies which specifically target the intestinal epithelium. Given this context, dietary components and their microbial metabolites offer promising potential as novel therapeutic targets for enhancing intestinal epithelial integrity.

Despite this potential, a significant gap remains in our understanding of which dietary components can effectively reduce the risk of relapse and help maintain remission in UC patients. The recent American Gastroenterology Association (AGA) Clinical Practice Update on diet and nutritional therapies in patients with IBD recommended all patients to follow a Mediterranean diet based on the evidence demonstrating its anti-inflammatory effects in IBD[6]. Adhering to a healthy, balanced Mediterranean-style diet enriched in fresh fruits, vegetables, whole grains, beans, and olive oil[7] may reduce the risk of flare in UC patients[8]. However, it remains unclear whether the minimized dietary components, such as red and processed meat, contribute to its efficacy or whether the enriched components and their microbial metabolites are more important. Overall, expanding our knowledge of how dietary components and their metabolism by intestinal bacteria influence intestinal epithelial integrity is essential for understanding the pathophysiology of UC and developing targeted therapies for IECs.

In this study, we investigate gut microbial metabolites that can enhance intestinal epithelial integrity by decreasing IEC permeability, clarify their metabolic status in UC patients, and examine their anti-inflammatory effects using a mouse model of colitis.

## Materials and Methods

### Mice

Female C57BL/6 mice were purchased from Oriental Yeast India Pvt. Ltd. All mice were housed and fed in a dedicated pathogen-free facility and maintained at standard laboratory conditions throughout the experiment. The animal experiments were conducted according to the protocol reviewed and approved by the Institutional Animal Care and Use Committee of the Jikei University School of Medicine (approval number: 2021-085).

### *Induction of colitis by dextran sodium sulfat*e

Experimental colitis was induced in 8-week-old female mice by administering 3% w/v dextran sodium sulfate (DSS) in drinking water for 4 days, followed by a regime of 3 days of normal water[9]. In the current study, the treatment group received intrarectal or oral administration of a 5mM solution of 3-aminobenzoic acid (3-ABA) on day 1, day 3 and day 5. In contrast, the control group received rectal or oral administration of PBS on the same days. On day 8, mice were euthanized using 5% isoflurane and subsequently killed by cervical dislocation. The colon was removed, measured for the length, and used for the subsequent analysis.

### Microscopic assessment of colitis

Transverse sections were prepared from rectums, fixed in 10% formaldehyde, and embedded in paraffin for histological analysis. Sections were stained with hematoxylin and eosin (HE), and histological damage was quantified as we previously reported[10] in a blinded manner.

### Human Subjects, Samples, and Clinical Information

Fecal samples were obtained from patients with an established diagnosis of UC and endoscopically active inflammation in the colon and healthy non-IBD (NIBD) controls between April 2020 and January 2024. Cross-sectional clinical information was collected at the time of sampling and summarized in **Supplementary Table 1**. All patients with UC were taking 5-ASA. Endoscopic disease activity of UC was evaluated by Mayo Endoscopic Subscore (MES) with active inflammation defined as MES of 2 or 3[11].

### Metabolomic Analysis

Samples were homogenized and subjected to the extraction of aminobenzoic acid (ABA) using methanol. Identification of structural isomers of ABA, including 2-ABA, 3-ABA, and 4-ABA, was conducted using liquid chromatography coupled with quadrupole time-of-flight mass spectrometry (LC-QTOF-MS). The separation was achieved on an InterSustain C18 column (GL Sciences Inc., Tokyo, Japan) with a suitable mobile phase. The analysis was performed on a Maxis Impact II (Bruker Daltonics, Bremen, Germany) with positive electrospray ionization (ESI)[12]. The fragment pattern of 3-ABA was further confirmed in fecal samples using liquid chromatography-tandem mass spectrometry (LC-MS/MS).

### DNA extraction and next-generation sequencing of the fecal bacterial 16S rRNA genes

Bacterial DNA was extracted from human fecal samples using phenol-chloroform extraction method with glass beads and purified using a High Pure PCR Template Preparation Kit (Roche Diagnostics, Mannheim, Germany) according to the manufacturer’s instructions. The variable V3-4 regions of the 16S rRNA gene were amplified by PCR using the universal 16S primer sets 341F and 805R which contain the Illumina index and sequencing adapter overhangs. The amplicons were purified using SPRIselect (Beckman Coulter Inc., CA, USA). The DNA was quantified using a QuantiFluor One dsDNA system (Promega Corporation, WI, USA) and sequenced using a MiSeq Reagent Kit v3 and MiSeq sequencer (Illumina Inc., CA, USA)[13]. The obtained data was analyzed using Quantitative Insights Into Microbial Ecology (QIIME2) version 2019.10 followed by A Linear Discriminant Analysis (LDA) Effect Size (LEfSe) analysis to understand the difference in the microbial community between UC patients and NIBD controls[14]. Bacterial taxa with LDA scores of > 2.0 and a P value of < 0.05 were considered significantly enriched. Alpha diversity was analyzed using the Shannon index, while beta diversity was analyzed using unweighted and weighted UniFrac distances with statistical significance determined by the Permutational Multivariate Analysis of Variance (PERMANOVA) test. The relationship between luminal 3-ABA and the relative abundance of intestinal bacteria was analyzed using Spearman’s correlation coefficient.

### Predictive Functional Profiling of Gut Microbial Communities

The metagenomics-based function of the microbiome was estimated using the Phylogenetic Investigation of Communities by Reconstruction of Unobserved States (PICRUSt) v2.4.1 to obtain relative Kyoto Encyclopedia of Genes and Genomes (KEGG) pathway abundance information[15]. Pathways with LDA scores of > 2.0 and a P value of < 0.05 were considered significantly enriched. The predicted data were collapsed into hierarchical categories, and the relative abundances of gut metabolic functions were calculated and graphed using R software Version 4.1.0.

### Cell culture

The human colonic epithelial cell line Caco2 (ATCC; HTB-37) was cultured in complete Eagle’s Minimum Essential Medium (EMEM) containing 10% heat-inactivated fetal bovine serum, 100 mg/ml streptomycin, and 100 mg/ml penicillin, and incubated in a 5% CO2 incubator at 37 °C.

### Epithelial Permeability Assay

Caco2 cells were cultured on HTS Transwell-96 Tissue Culture Systems (Corning Incorporated, Corning, NY, USA) to form cell monolayers and stimulated with a library of 119 gut microbial metabolites (MedchemExpress, NJ, USA) for 3 days at a final concentration of 10 μM unless indicated otherwise. The electrical resistance of cell monolayers was measured on day 4 (before stimulation) and day 7 (after 3 days of stimulation) using an EVOM2 epithelial Volt/ Ohm meter (World Precision Instruments, FL, USA). The resistance of cell monolayers was corrected by subtracting the resistance of wells without cells and normalized by multiplying by the effective well surface area to provide transepithelial electrical resistance (TEER) in the unit of Ω cm^2^. Changes in TEER between day 4 and day 7 were evaluated by calculating ΔTEER% as we previously reported[16]. The same experiment was repeated on 12-well cell culture inserts (BD Falcon, CA, USA). On day 4, fluorescein isothiocyanate (FITC)-dextran (70 kDa, Sigma-Aldrich, MO, USA) was added to the culture inserts at a final concentration of 10 μM. The concentration of FITC-dextran permeated to the lower chamber was measured using a Qubit 4 Fluorometer (Invitrogen, CA, USA). Intestinal epithelial permeability was also evaluated in mice by measuring the levels of FITC-dextran in the serum after oral gavage of FITC-dextran for 3 hours.

### Reverse-Transcriptase qPCR Analysis

Total RNA was extracted from the rectum of mice and purified with the Single Cell RNA Purification Kit (Norgen Biotek, ON, Canada) according to the manufacturer’s instructions. Complementary DNA was synthesized from 100 ng of RNA using the High-Capacity Complementary DNA Reverse Transcription Kit (Thermo Fisher Scientific, MA, USA). qPCR for mRNAs was performed on the QuantStudio 3 RT-PCR system using PowerUp SYBR Green Master Mix (Thermo Fisher Science) and primers listed in **Supplementary Table 2**. Gene expression was normalized to *GAPDH* or the 16S rRNA V4 sequences for *mabA* and *mabB*. Relative changes in gene expression were compared across samples using the 2^-ΔΔCT^ method[16].

### RNA-Sequencing

Total RNA was extracted from Caco2 cells after stimulation with 5mM of 3-ABA for 24 hours. RNA integrity was evaluated using an Agilent 2100 Bioanalyzer and confirmed to have an RNA Integrity Number (RIN) of 7 or above. RNA-seq libraries were prepared using the Illumina SMARTer Stranded RNA-Seq Kit. Paired-end sequencing was performed on the Illumina NovaSeq 6000 platform (Gene Expression Omnibus [GEO] accession no. GSE270624) and analyzed using RaNA-Seq and RNAseqChef, cloud platforms for the analysis and visualization of RNA-Seq data[17, 18]. Gene expression profiles were compared between 3-ABA-stimulated and non-stimulated cells (N=3 per group).

### Statistical Analysis

All numeric data are expressed as means ± standard deviation or median ± interquartile range (IQR). A Mann–Whitney test analyzed differences between 2 groups. Differences among 3 groups were analyzed by a Kruskal–Wallis test followed by Dunn’s multiple comparison test. P values less than 0.05 were considered significant. GraphPad Prism version 9.5.1 (GraphPad Software, CA, USA) was used for these data analyses. A differential expression analysis of RNA-seq was performed using DESeq2[19], with FDR-adjusted P values less than 0.05 being considered statistically significant.

### *Ethical Statemen*t

This study was conducted in accordance with the Declaration of Helsinki and Good Clinical Practice guidelines. The study protocol was approved by the institutional review board of the Jikei University School of Medicine (approval number: 32-164-10245). All participants provided written informed consent before inclusion in the study. All participants were identified by number and not by name or any protected health information.

## Results

### Aminobenzoic acid decreased epithelial permeability in Caco2 cells

To explore gut microbial metabolites that can enhance intestinal epithelial integrity, we stimulated Caco2 cells with a library of 119 gut microbial metabolites and calculated the percentage changes of TEER (ΔTEER%) in vitro (**Supplementary Table 3**). Among 10 metabolites with the highest ΔTEER%, 3 were tryptophan metabolites including tryptophol, tryptamine, and N-acetyl-L-tryptophan. The others included L-lysine hydrochloride, L-Ascoribic acid, Rhamnose monohydrate, 4-aminobenzoic acid (4-ABA), deoxycholic acid, and spermidine. Notably, 4-ABA exhibited the largest increase in TEER and decreased IEC permeability in repeated experiments (**Supplementary Figure 1**). Further experiments with 2-ABA and 3-ABA, two structural isomers of 4-ABA in the natural environment, found the largest elevation of TEER after stimulation with 3-ABA at a dose of 5 mM (median ΔTEER%, control vs. 3-ABA, 18.2 vs. 144.2 %, p<0.05, **Figure 1A**). 5-ASA also elevated TEER in Caco2 cells, but this effect size was numerically lower than 3-ABA. 3-ABA increased TEER in a dose-dependent manner and significantly reduced the permeation of FITC-dextran across Caco2 cell monolayers (**Figure 1B**).

**Figure 1.**
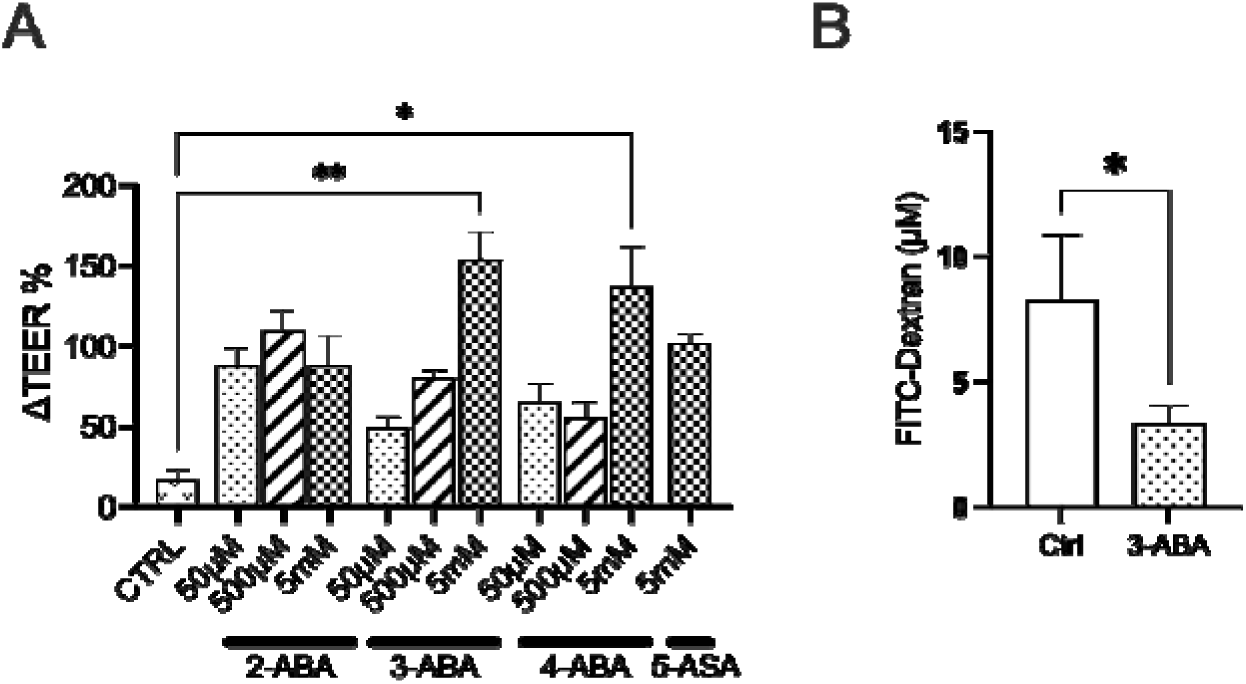
3-aminobenzoic acid reduced epithelial permeability in Caco2 cell monolayers. (A) Dose-dependent changes of ΔTEER% in Caco2 cells stimulated with 2-, 3-, and 4-aminobenzoic acid (ABA). (B) The concentration of FITC-dextran permeated across Caco2 cell monolayers stimulated with 5mM 3-ABA. *P<0.05, **P<0.01 compared to the non-stimulated controls. N=3-4 per group.

### 3-aminobenzoic acid was abundant in beans and decreased in the feces of patients with UC

To understand the source and metabolism of 3-ABA within the human intestine, we first investigated its presence in the diet using the Food Metabolome Repository[20]. A precursor search based on the mass-to-charge ratio estimated the presence of 3-ABA with the highest probability in beans, such as azuki beans, soybeans, and chickpeas (**Figure 2A**). We substantiated the presence of 3-ABA in these beans through LC-QTOF-MS analysis (**Supplementary Figure 2A**).

**Figure 2.**
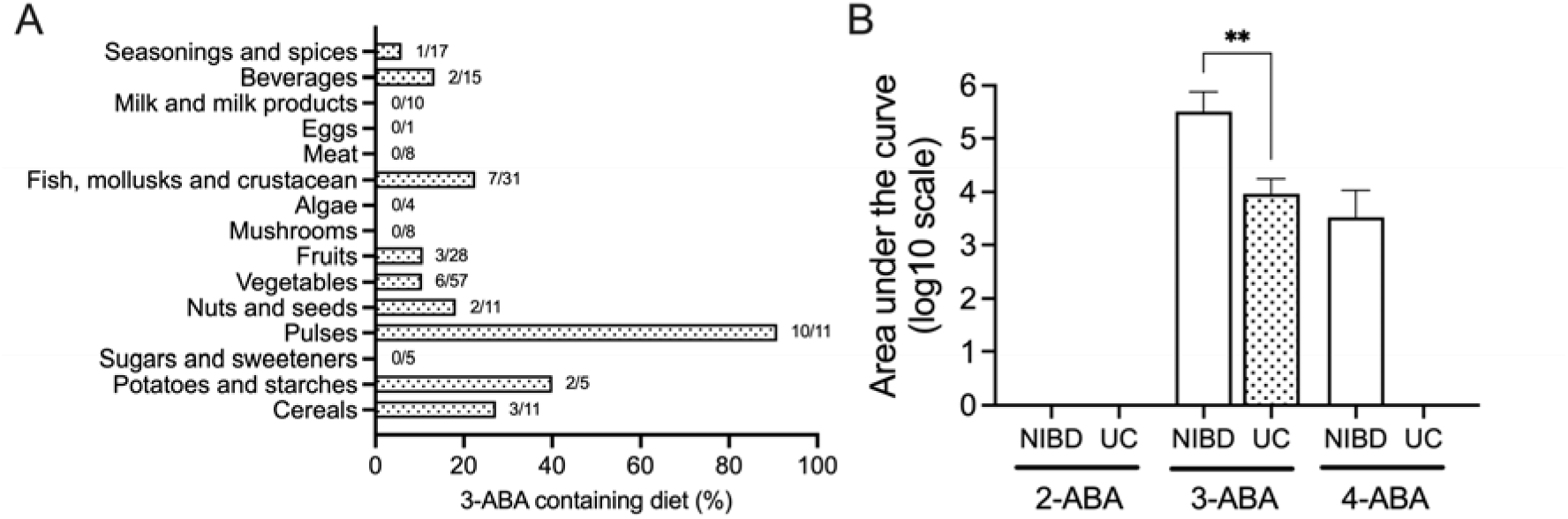
3-aminobenzoic acid was abundant in beans and decreased in the feces of patients with ulcerative colitis (UC). (A) The detection rate of 3-aminobenzoic acid (3-ABA) in various types of diet registered in the Food Metabolome Repository. The number of diets estimated to contain 3-ABA is shown as "number estimated/total registered" above the bars for each dietary category. (B) The abundance of ABA in the fecal samples of patients with UC and healthy non-inflammatory bowel disease (NIBD) controls. **P<0.01. N=5 per group.

Next, we confirmed 3-ABA in the intestines of healthy NIBD individuals comparing its levels with those of the other structural isomers. We revealed the dominant presence of 3-ABA in the fecal samples of NIBD individuals, while detecting minimal presences of 2-ABA or 4-ABA (**Figure 2B**). The fecal amount of 3-ABA was significantly lower in UC patients than in NIBD controls (**Figure 2B** and **Supplementary Figure 2B**).

### Degradation of luminal 3-aminobenzoic acid was accelerated in patients with UC

We hypothesized that dysbiosis alters gut microbial functions and accelerates the degradation of 3-ABA in the intestine of UC patients. To address this hypothesis, we conducted a 16S rRNA metagenomic analysis and predicted the functions of the gut microbiota. The metagenomic analysis revealed a significantly reduced diversity of gut microbiota in UC patients compared to NIBD (**Supplementary Figure 3A**). A LEfSe analysis identified a decreased abundance of *Lachnospiraceae* and an increased abundance of *Pseudomonadota* (*Escherichia_Shigella*) and Bacillota (*Negativibacillus* and *Granulicatella*) in patients with UC compared to NIBD controls (**Supplementary Figure 3B**). These findings substantiated the presence of dysbiosis in the UC patients who participated in this study. Functional prediction analysis demonstrated that UC patients were significantly more enriched with the bacteria involved in the degradation of ABA than NIBD controls (**Figure 3A**). Relative expression levels of *mabA*, the gene encoding a bacterial enzyme involved in the degradation of 3-ABA into 5-ASA[21], were significantly higher in UC patients than in NIBD controls (**Figure 3B**). In contrast, there was no significant difference in the expression of *mabB*, the gene encoding a bacterial enzyme involved in the degradation of 5-ASA into 4-amino-6-oxohepta-2,4-dienedioic acid[21], between UC and NIBD controls (**Supplementary Figure 3C**). LC-QTOF-MS analysis identified 5-ASA and its major microbial metabolites, such as N-acetyl-5-ASA, N-Propionyl-5-ASA, and N-Butyryl-5-ASA[22], but did not find 4-amino-6-oxohepta-2,4-dienedioic acid in the fecal samples of UC patients (**Figure 3C**). Correlation analysis identified a significant inverse relationship between luminal 3-ABA and the relative abundance of *Escherichia_Shigella* (r = - 0.70, **Supplementary figure 3D**).

**Figure 3.**
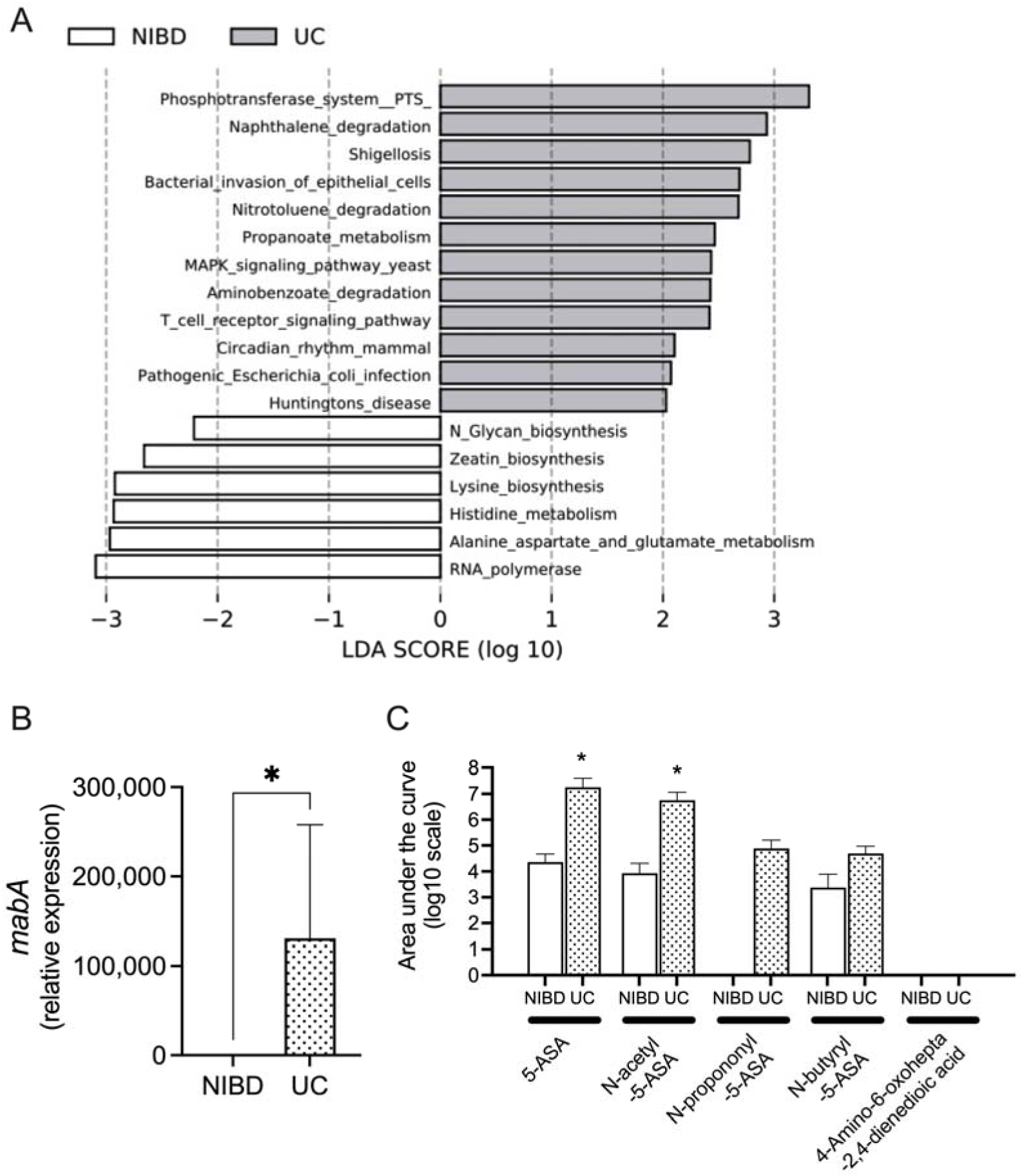
The degradation of luminal 3-aminobenzoic acid was accelerated in patients with ulcerative colitis (UC). (A) Functional prediction analysis of the gut microbiota in patients with UC and healthy non-inflammatory bowel disease (NIBD) controls. ( B ) The expression levels of *mabA* in the fecal DNA. (C) The relative abundance of 5-ASA and its microbial metabolites in the fecal samples of patients with UC and NIBD controls. *P<0.05 compared to NIBD. N=5-7 per group.

### 3-aminobenzoic acid regulated tight junction in Caco2 cells

To clarify the mechanism through which 3-ABA reduces IEC permeability, we conducted a transcriptional analysis on Caco2 cells stimulated with 3-ABA. Gene enrichment analysis using the Kyoto Encyclopedia of Genes and Genomes (KEGG) pathway revealed that differentially expressed genes (DEGs) in 3-ABA-stimulated cells are significantly enriched in the tight junction regulatory pathway (**Table 1**). Notably, the expression of *CLDN1* and *Tight Junction Protein 1* (*TJP1*), which encode the tight junction proteins, Claudin1 and Zonula occludens-1 (ZO-1), respectively, were significantly elevated in the 3-ABA-stimulated cells compared to the non-stimulated controls (**Supplementary Figure 4A**). In addition, gene enrichment analysis identified DEGs significantly enriched in the PPAR signaling pathway, which regulates *CLDN1*[23] and *TJP1*[24], in the 3-ABA-stimulated cells. The expression of genes downstream of PPARγ, such as *CPT1A*, *SORBS1*, and *ACSL5* was significantly higher in the 3-ABA-stimulated cells than in controls (**Supplementary Figure 4B**).

**Table 1.** Gene Set Enrichment Analysis of differentially expressed genes in Caco2 cells stimulated with 3-aminobenzoic acid compared with non-stimulated cells.

| Description | Gene Set ID | DEG Count | Gene Ratio<br>(DEG/total gene) | Adjusted p-value |
| --- | --- | --- | --- | --- |
| Pathways in cancer | M12868 | 314 | 0.068 | 0.00000026 |
| MAPK signaling pathway | M10792 | 254 | 0.055 | 0.00024755 |
| Regulation of actin cytoskeleton | M18306 | 199 | 0.043 | 0.01162371 |
| Focal adhesion | M7253 | 192 | 0.042 | 0.00017266 |
| Endocytosis | M1519 | 180 | 0.039 | 0.00000009 |
| Purine metabolism | M14314 | 151 | 0.033 | 0.00515517 |
| Wnt signaling pathway | M19428 | 142 | 0.031 | 0.01986812 |
| Ubiquitin mediated proteolysis | M15247 | 133 | 0.029 | 0.00008051 |
| Insulin signaling pathway | M18155 | 131 | 0.029 | 0.00479758 |
| Spliceosome | M2044 | 126 | 0.027 | 0.00003655 |
| Axon guidance | M5539 | 126 | 0.027 | 0.00055933 |
| Neurotrophin signaling pathway | M16763 | 124 | 0.027 | 0.00017266 |
| Tight junction | M11355 | 124 | 0.027 | 0.03086545 |
| Cell cycle | M7963 | 123 | 0.027 | 0.00017266 |

DEG, differentially expressed gene

### 3-aminobenzoic acid attenuated DSS-induced colitis

The epithelial barrier-enhancing effect of 3-ABA and its potential conversion to 5-ASA in the inflamed colon implied its efficacy for preventing and improving intestinal inflammation in UC patients. Therefore, we examined the anti-inflammatory effect of 3-ABA in DSS-induced colitis mice. Rectal administration of 3-ABA significantly improved body weight loss and prevented the shortening of colon length (**Figure 4A**, **Figure 4B**, and **Supplementary Figure 5A**). The histologic scores were significantly lower in the 3-ABA-treated mice than in the controls. (**Figure 4C** and **Supplementary Figure 5B**). The expression of pro-inflammatory cytokines including tumor necrosis factor (TNF), interleukin-1b (Il-1b), and Il-6 was decreased in the colon of 3-ABA-treated mice compared to the control mice (**Figure 4D**). The serum concentration of the FITC-dextran permeated across the intestinal epithelium was numerically lower in the 3-ABA-treated mice than in the controls (**Figure 4E**). Oral administration of 3-ABA showed similar but less dramatic effects with numerically improved body weight loss, colon length shortening, histological score, and the expression of pro-inflammatory cytokines in the colon (**Supplementary Figure 6A-D**).

**Figure 4.**
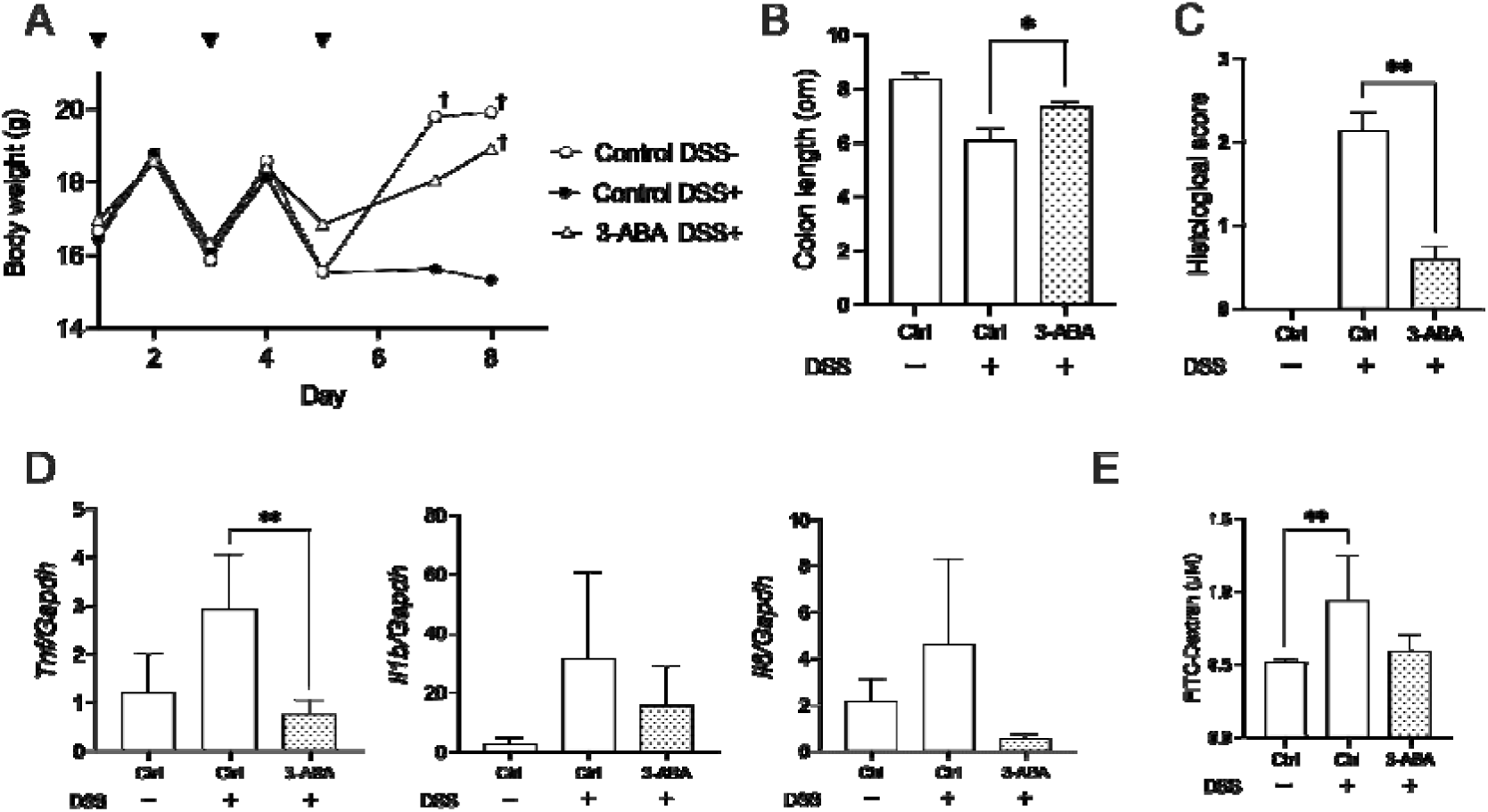
Rectal administration of 3-aminobenzoic acid attenuated DSS-induced colitis in mice. (A) Body weight change during the experiment. Arrowheads indicate the timing of rectal drug administration. ^†^P<0.05 compared to the DSS-treated control mice (Control DSS+). The colon length (B), histologic score (C), gene expression of inflammatory cytokines (D), and serum levels of FITC-Dextran (E) were evaluated on day 8. *P<0.05 **P<0.01 N=6 per group.

## Discussion

3-ABA, also known as meta-aminobenzoic acid, is an organic compound commonly used as a fundamental component in the synthesis of various pharmaceuticals, dyes, and other organic compounds[25]. Unlike its structural isomers, 2-ABA and 4-ABA, which are synthesized and utilized by intestinal bacteria as an essential substrate for synthesizing folic acid[26], the biological function and metabolism of 3-ABA in the intestinal tract remains largely unexplored. In this study, we demonstrated that 3-ABA was abundant in beans, enhanced intestinal epithelial integrity by reducing IEC permeability in vitro, and exhibited an anti-inflammatory effect in a mouse model of colitis. The degradation of luminal 3-ABA was accelerated in UC patients and potentially metabolized into 5-ASA via increased mabA expression in intestinal bacteria.

Recent studies have highlighted the importance of gut microbiota for maintaining intestinal epithelial integrity. For example, germ-free mice exhibited enhanced maturation of colonic barrier structure and increased resistance to chemically induced colonic injury after colonization with human gut microbiota[27]. Although the precise mechanism by which microbiota regulate colonic epithelial integrity is not fully understood, it is partially attributed to their metabolism of dietary components, including short-chain fatty acids, tryptophan, and polyamines[28]. These metabolites regulate IEC permeability by modulating the expression of junctional proteins and help maintain intestinal epithelial homeostasis in health. In this study, we observed elevations of TEER in Caco2 cells stimulated with several tryptophan metabolites and spermidine, one of the polyamines, confirming their roles in maintaining intestinal epithelial integrity. Tryptophan metabolites, such as indole-3-ethanol, indole-3-pyruvate, and indole-3-aldehyde have been shown to alleviate DSS-induced colitis in mice by preserving the integrity of the apical junctional complex and its associated actin regulatory proteins, thereby protecting against increased IEC permeability[29]. Oral administration of spermidine also alleviated DSS-induced colitis and prevented the development of colitis-associated dysplasia in mice[30]. In contrast, the role of ABA in maintaining intestinal epithelial homeostasis has not been previously investigated. Reduced IEC permeability in ABA-stimulated Caco2 cells in this study suggested the contribution of ABA in regulating intestinal epithelial integrity. The relative abundance of 3-ABA among three structural isomers in the human intestine and the remarkable improvement of DSS-induced colitis in mice by rectal 3-ABA administration further emphasized the importance of this structural isomer in the colon.

Notably, 3-ABA was contained in the diet, especially in beans, and potentially converted into 5-ASA, an established therapeutic molecule for UC, by mabA in the inflamed colon of UC patients. The significant inverse relationship between luminal 3-ABA and the abundance of *Escherichia_Shigella*, which was enriched in UC, implied its involvement in the 3-ABA degradation. Since the specific bacteria expressing *mabA* in the intestine are currently unknown, further investigation using shotgun metagenomic sequencing will be necessary to identify those involved in the mabA-dependent degradation of 3-ABA. In contrast to *mabA*, UC patients did not show a significant up-regulation of *mabB* in their feces. The absence of its metabolic product in the feces of UC patients suggested that the degradation of 5-ASA into 4-amino-6-oxohepta-2,4-dienedioic acid by mabB is not the dominant metabolic pathway of 5-ASA in the inflamed colon. Previous studies reported the improvement of DSS-induced colitis in mice through the consumption of kidney beans and azuki beans[31, 32], although the mechanism remains unclear. In addition, recent studies have shown that the Mediterranean diet, which is characterized by a high abundance of beans[7], contributes to reducing intestinal inflammation in UC patients[33, 34]. These findings imply that proactive intake of 3-ABA or a 3-ABA-rich diet could attenuate colonic inflammation in UC patients regardless of the accelerated degradation of 3-ABA by intestinal bacteria.

Given the current reliance on immune-suppressive drugs for UC treatment, 3-ABA might offer a novel therapeutic approach that avoids the risks of infections and malignancies associated with immune suppression. Moreover, combining 3-ABA with other drugs might enhance therapeutic efficacy in patients suffering from UC resistant to existing treatments. The greater elevation of TEER in the 3-ABA-stimulated cells compared to the 5-ASA-stimulated cells in this study implied its superior therapeutic potential for UC. Although the mechanisms by which 3-ABA reduces IEC permeability and attenuates mucosal inflammation remain unclear, we found upregulated *CLDN1* and *TJP1* expression in 3-ABA-stimulated Caco2 cells. Concomitant upregulation of the genes downstream of PPARγ implied the involvement of PPARγ activation in regulating these genes as previously reported[23, 24]. As the absence of *CLDN1* and *TJP1* increases epithelial permeability[35, 36] and the lack of *TJP1* in IEC further disrupts mucosal repair in mice by attenuating Wnt-β-catenin signaling and inducing abortive IEC proliferation[36], 3-ABA might preserve intestinal epithelial homeostasis by inducing these junctional proteins in IECs through activation of PPARγ.

There are several limitations in this study. First, we did not assess the fecal levels of 3-ABA in UC patients who achieved endoscopic remission. Consequently, it remains plausible that alterations in the gut microbiota driven by inflammation could accelerate 3-ABA degradation. Patients in remission might have higher luminal 3-ABA levels like healthy individuals than those with active disease. Investigating the correlation between luminal 3-ABA and disease activity could shed light on this question and may further reveal the potential utility of measuring fecal 3-ABA for monitoring disease activity or predicting relapse in patients with UC. Second, we collected fecal samples from a small number of patients with no dietary information. Given that 3-ABA is contained in the diet, particularly enriched in beans, dietary information is necessary to assess its impact on luminal 3-ABA levels. Future studies incorporating a larger number of patients with comprehensive dietary information would help validate our findings and determine the amount of dietary 3-ABA necessary to prevent or attenuate colitis in UC patients. Third, we did not demonstrate the direct conversion of 3-ABA into 5-ASA by mabA in the human colon. Since all patients with UC in this study were taking 5-ASA as the first-line drug, it was difficult to identify 5-ASA derived from 3-ABA in their fecal samples. Finally, we did not explore the effect of 3-ABA on non-IEC cells. Considering that PPARγ is expressed in immune cells, such as T lymphocytes and macrophages, and regulates their functions[37, 38], 3-ABA might also mitigate colonic inflammation by activating PPARγ in these immune cells. Understanding the effect of 3-ABA on immune cell functions through PPARγ could provide further insights into its role in maintaining intestinal homeostasis and potentially its efficacy in the treatment of UC.

Despite these limitations, our present study is significant because we identified 3-ABA as a novel dietary component regulating intestinal epithelial integrity and elucidated its metabolic difference between health and disease. The anti-inflammatory effect of 3-ABA and its potential conversion into 5-ASA in the inflamed colon indicated its promising efficacy in the treatment of UC. Further understanding of the role of 3-ABA in maintaining intestinal homeostasis and the pathophysiology of UC will help pave the concrete path toward the development of novel therapeutic strategies using 3-ABA.

## Supporting information

Supplemental Files.docx

## Acknowledgements

None

## Disclosure of funding

This work was supported by the Japanese Society for the Promotion of Science “KAKENHI” grant number 21K15985.

## Conflicts of Interest

T. Toyonaga received research grants from Pfizer Inc. M. Saruta receives lecture fees from Janssen Pharma K.K., Takeda Pharmaceutical Co.,Ltd., Mitsubishi Tanabe Pharma Co., Ltd., and EA Pharma Co., Ltd., advisory/consulting board fees from EA Pharma Co., Ltd., and grants from Zeria Pharmaceutical Co., Ltd., Mochida Pharmaceutical Co.,Ltd., EA Pharma Co., and Ltd. EP-CRSU Co., Ltd.

## Contributions

Takahiko Toyonaga devised the conceptual ideas of the study. Miho Tanaka, Takahiko Toyonaga, Fumiyuki Nakagawa, Takeo Iwamoto, Akira Komatsu, Natsuki Sumiyoshi, Naoki Shibuya, Ayaka Minemura, Tadashi Ariyoshi, Asami Matsumoto, Kentaro Oka, Masayuki Shimoda contribute to the acquisition of data, and analysis and interpretation of data. Miho Tanaka and Takahiko Toyonaga also contributed to drafting the article. Masayuki Saruta supervised the project. All authors had full access to all the data in the study, revised the article critically for important intellectual content, and had final responsibility for the decision to submit for publication.

## Reference

1 Lavelle A, Sokol H. Gut microbiota-derived metabolites as key actors in inflammatory bowel disease. Nat Rev Gastroenterol Hepatol 2020;17:223–37.

2 Lee D, Albenberg L, Compher C, Baldassano R, Piccoli D, Lewis JD, et al. Diet in the pathogenesis and treatment of inflammatory bowel diseases. Gastroenterology 2015;148:1087–106.

3 Hou JK, Abraham B, El-Serag H. Dietary intake and risk of developing inflammatory bowel disease: a systematic review of the literature. Am J Gastroenterol 2011;106:563–73.

4 Kobayashi T, Siegmund B, Le Berre C, Wei SC, Ferrante M, Shen B, et al. Ulcerative colitis. Nat Rev Dis Primers 2020;6:74.

5 Sehgal P, Colombel JF, Aboubakr A, Narula N. Systematic review: safety of mesalazine in ulcerative colitis. Aliment Pharmacol Ther 2018;47:1597–609.

6 Hashash JG, Elkins J, Lewis JD, Binion DG. AGA Clinical Practice Update on Diet and Nutritional Therapies in Patients With Inflammatory Bowel Disease: Expert Review. Gastroenterology 2024;166:521–32.

7 Yannakoulia M, Scarmeas N. Diets. N Engl J Med 2024;390:2098–106.

8 Chicco F, Magri S, Cingolani A, Paduano D, Pesenti M, Zara F, et al. Multidimensional Impact of Mediterranean Diet on IBD Patients. Inflamm Bowel Dis 2021;27:1–9.

9 Jeengar MK, Thummuri D, Magnusson M, Naidu VGM, Uppugunduri S. Uridine Ameliorates Dextran Sulfate Sodium (DSS)-Induced Colitis in Mice. Sci Rep 2017;7:3924.

10 Egger B, Procaccino F, Lakshmanan J, Reinshagen M, Hoffmann P, Patel A, et al. Mice lacking transforming growth factor alpha have an increased susceptibility to dextran sulfate-induced colitis. Gastroenterology 1997;113:825–32.

11 Turner D, Ricciuto A, Lewis A, D’Amico F, Dhaliwal J, Griffiths AM, et al. STRIDE-II: An Update on the Selecting Therapeutic Targets in Inflammatory Bowel Disease (STRIDE) Initiative of the International Organization for the Study of IBD (IOIBD): Determining Therapeutic Goals for Treat-to-Target strategies in IBD. Gastroenterology 2021;160:1570–83.

12 Wu C, Iwamoto T, Hossain MA, Akiyama K, Igarashi J, Miyajima T, et al. A combination of 7-ketocholesterol, lysosphingomyelin and bile acid-408 to diagnose Niemann-Pick disease type C using LC-MS/MS. PLoS One 2020;15:e0238624.

13 Kusamoto A, Harada M, Minemura A, Matsumoto A, Oka K, Takahashi M, et al. Effects of the prenatal and postnatal nurturing environment on the phenotype and gut microbiota of mice with polycystic ovary syndrome induced by prenatal androgen exposure: a cross-fostering study. Front Cell Dev Biol 2024;12:1365624.

14 Bolyen E, Rideout JR, Dillon MR, Bokulich NA, Abnet CC, Al-Ghalith GA, et al. Reproducible, interactive, scalable and extensible microbiome data science using QIIME 2. Nat Biotechnol 2019;37:852–7.

15 Ariyoshi T, Hagihara M, Tomono S, Eguchi S, Minemura A, Miura D, et al. Clostridium butyricum MIYAIRI 588 Modifies Bacterial Composition under Antibiotic-Induced Dysbiosis for the Activation of Interactions via Lipid Metabolism between the Gut Microbiome and the Host. Biomedicines 2021;9.

16 Toyonaga T, Steinbach EC, Keith BP, Barrow JB, Schaner MR, Wolber EA, et al. Decreased Colonic Activin Receptor-Like Kinase 1 Disrupts Epithelial Barrier Integrity in Patients With Crohn’s Disease. Cell Mol Gastroenterol Hepatol 2020;10:779–96.

17 Prieto C, Barrios D. RaNA-Seq: Interactive RNA-Seq analysis from FASTQ files to functional analysis. Bioinformatics 2019.

18 Etoh K, Nakao M. A web-based integrative transcriptome analysis, RNAseqChef, uncovers the cell/tissue type-dependent action of sulforaphane. J Biol Chem 2023;299:104810.

19 Love MI, Huber W, Anders S. Moderated estimation of fold change and dispersion for RNA-seq data with DESeq2. Genome Biol 2014;15:550.

20 Sakurai N, Shibata D. Tools and databases for an integrated metabolite annotation environment for liquid chromatography-mass spectrometry-based untargeted metabolomics. Carotenoid Science 2017;22:16–22.

21 Yu H, Zhao S, Lu W, Wang W, Guo L. A novel gene, encoding 3-aminobenzoate 6-monooxygenase, involved in 3-aminobenzoate degradation in Comamonas sp. strain QT12. Appl Microbiol Biotechnol 2018;102:4843–52.

22 Mehta RS, Mayers JR, Zhang Y, Bhosle A, Glasser NR, Nguyen LH, et al. Gut microbial metabolism of 5-ASA diminishes its clinical efficacy in inflammatory bowel disease. Nat Med 2023;29:700–9.

23 Ogasawara N, Kojima T, Go M, Ohkuni T, Koizumi J, Kamekura R, et al. PPARgamma agonists upregulate the barrier function of tight junctions via a PKC pathway in human nasal epithelial cells. Pharmacol Res 2010;61:489–98.

24 Huang Y, Wang C, Tian X, Mao Y, Hou B, Sun Y, et al. Pioglitazone Attenuates Experimental Colitis-Associated Hyperalgesia through Improving the Intestinal Barrier Dysfunction. Inflammation 2020;43:568–78.

25 Zhang H, Stephanopoulos G. Co-culture engineering for microbial biosynthesis of 3-amino-benzoic acid in Escherichia coli. Biotechnol J 2016;11:981–7.

26 Kratky M, Konecna K, Janousek J, Brablikova M, Jandourek O, Trejtnar F, et al. 4-Aminobenzoic Acid Derivatives: Converting Folate Precursor to Antimicrobial and Cytotoxic Agents. Biomolecules 2019;10.

27 Hayes CL, Dong J, Galipeau HJ, Jury J, McCarville J, Huang X, et al. Commensal microbiota induces colonic barrier structure and functions that contribute to homeostasis. Sci Rep 2018;8:14184.

28 Ghosh S, Whitley CS, Haribabu B, Jala VR. Regulation of Intestinal Barrier Function by Microbial Metabolites. Cell Mol Gastroenterol Hepatol 2021;11:1463–82.

29 Scott SA, Fu J, Chang PV. Microbial tryptophan metabolites regulate gut barrier function via the aryl hydrocarbon receptor. Proc Natl Acad Sci U S A 2020;117:19376–87.

30 Gobert AP, Latour YL, Asim M, Barry DP, Allaman MM, Finley JL, et al. Protective Role of Spermidine in Colitis and Colon Carcinogenesis. Gastroenterology 2022;162:813–27 e8.

31 Yook JS, Kim KA, Kim M, Cha YS. Black Adzuki Bean (Vigna angularis) Attenuates High-Fat Diet-Induced Colon Inflammation in Mice. J Med Food 2017;20:367–75.

32 Monk JM, Zhang CP, Wu W, Zarepoor L, Lu JT, Liu R, et al. White and dark kidney beans reduce colonic mucosal damage and inflammation in response to dextran sodium sulfate. J Nutr Biochem 2015;26:752–60.

33 Garcia-Mateo S, Martinez-Dominguez SJ, Gargallo-Puyuelo CJ, Gallego B, Alfambra E, Escuin M, et al. Healthy Lifestyle Is a Protective Factor from Moderate and Severe Relapses and Steroid Use in Inflammatory Bowel Disease: A Prospective Cohort Study. Inflamm Bowel Dis 2024.

34 Mathews SN, Lukin DJ. Shifting the Inflammatory Balance in Ulcerative Colitis Through Diet: A Mediterranean Diet Pattern is Associated with Improvements in Dysbiosis and Disease Activity. J Crohns Colitis 2023;17:1555–6.

35 Furuse M, Hata M, Furuse K, Yoshida Y, Haratake A, Sugitani Y, et al. Claudin-based tight junctions are crucial for the mammalian epidermal barrier: a lesson from claudin-1-deficient mice. J Cell Biol 2002;156:1099–111.

36 Kuo WT, Zuo L, Odenwald MA, Madha S, Singh G, Gurniak CB, et al. The Tight Junction Protein ZO-1 Is Dispensable for Barrier Function but Critical for Effective Mucosal Repair. Gastroenterology 2021;161:1924–39.

37 Guri AJ, Mohapatra SK, Horne WT, 2nd, Hontecillas R, Bassaganya-Riera J. The role of T cell PPAR gamma in mice with experimental inflammatory bowel disease. BMC Gastroenterol 2010;10:60.

38 Hontecillas R, Horne WT, Climent M, Guri AJ, Evans C, Zhang Y, et al. Immunoregulatory mechanisms of macrophage PPAR-gamma in mice with experimental inflammatory bowel disease. Mucosal Immunol 2011;4:304–13.

