## Supplemental Files.docx for "Exploring 3-Aminobenzoic Acid as a Therapeutic Dietary Component for Enhancing Intestinal Barrier Integrity in Ulcerative Colitis"

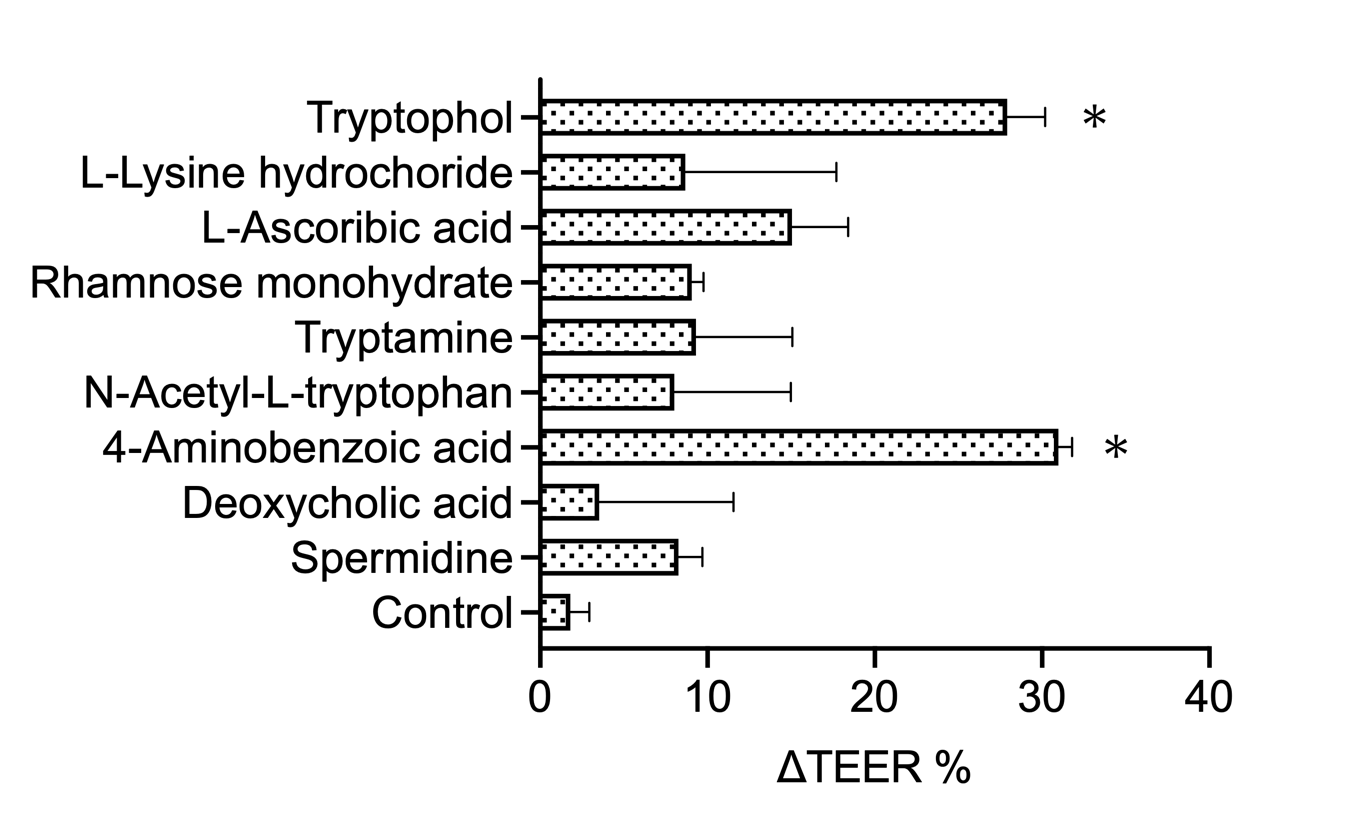

**Supplementary Figure 1.** The percentage changes of TEER (ΔTEER%) after stimulation with gut microbial metabolites in Caco2 cell monolayers. *P<0.05 compared to the non-stimulated controls. N=3 per group.

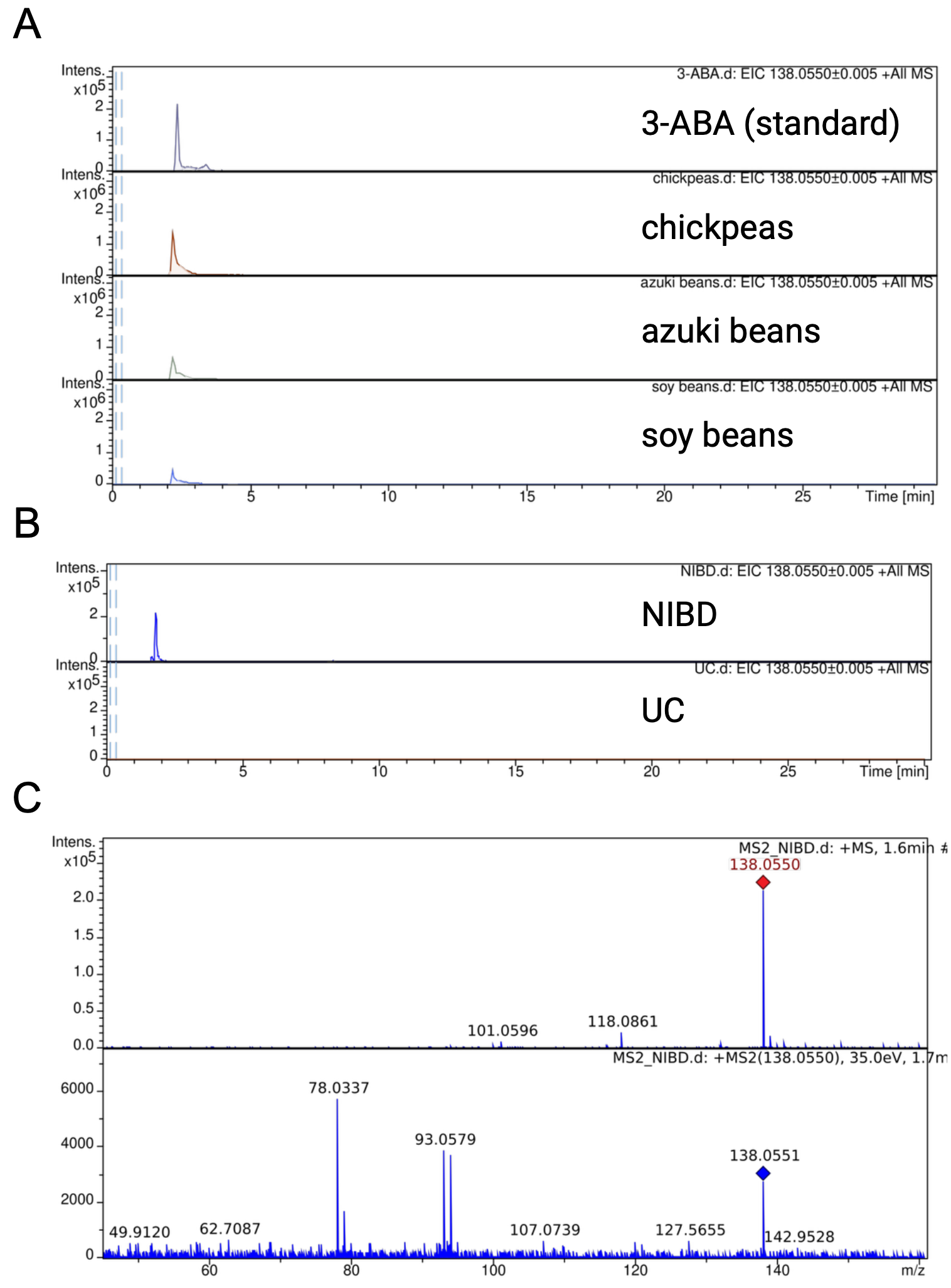

**Supplementary Figure 2.** (A) A representative image of LC-QTOF-MS targeted for 3-aminobenzoic acid (3-ABA) in various types of beans (chickpeas, azuki beans and soy beans) compared to the standard shown at the top of the figure. (B) A representative image of LC-QTOF-MS targeted for 3-ABA in patients with ulcerative colitis (UC) and healthy non-inflammatory bowel disease (NIBD) control. (C) A representative image of LC-MS/MS targeted for 3-ABA in NIBD controls.

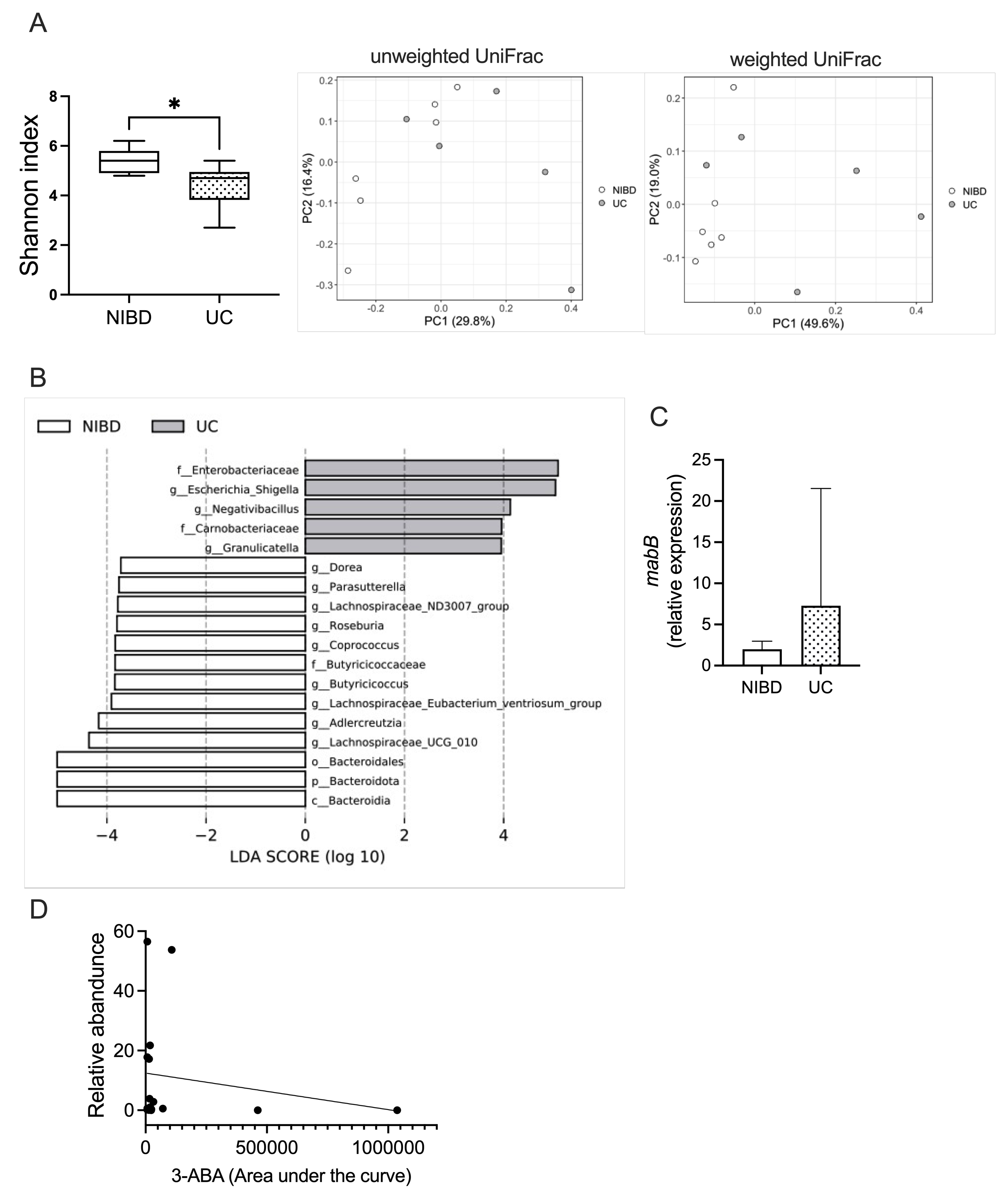

**Supplementary Figure 3.** The diversity (A) and composition (B) of the gut microbiota in patients with ulcerative colitis (UC) and healthy non-inflammatory bowel disease (NIBD) controls. (C) The expression levels of *mabB* in fecal DNA.

N=5-6 per group. (D) Correlation between luminal 3-ABA and the relative abundance of *Escherichia_Shigella* (r = - 0.70, p<0.05).

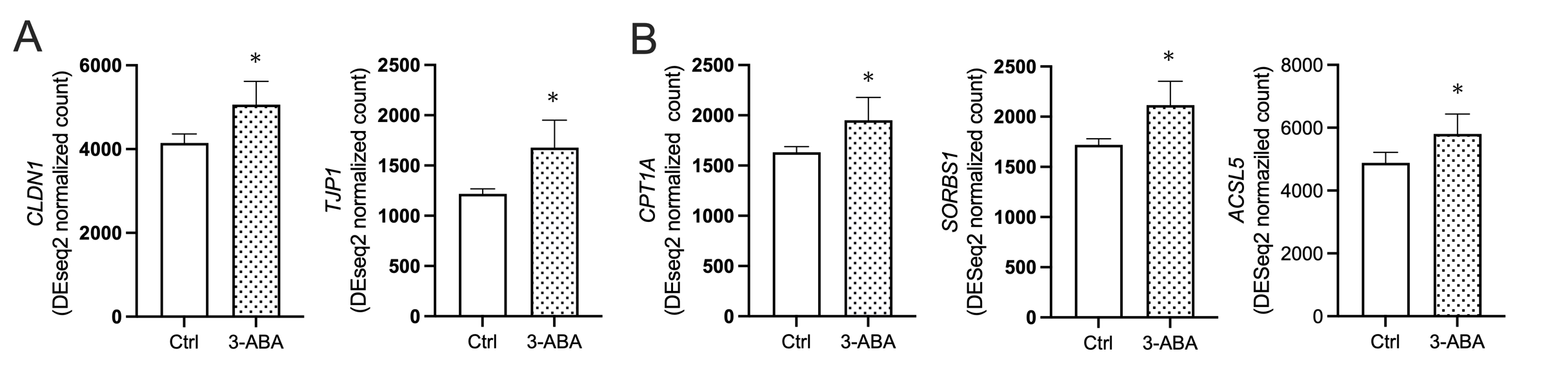

**Supplementary Figure 4.** Transcriptional analysis on the Caco2 cells stimulated with 3-aminobenzoic acid (3-ABA). (A) The expression of tight junctional molecules, including *CLDN1* and *TJP1*. (B) The expression of PPARγ downstream genes. *P<0.05 compared to the non-stimulated controls. N=3 per group.

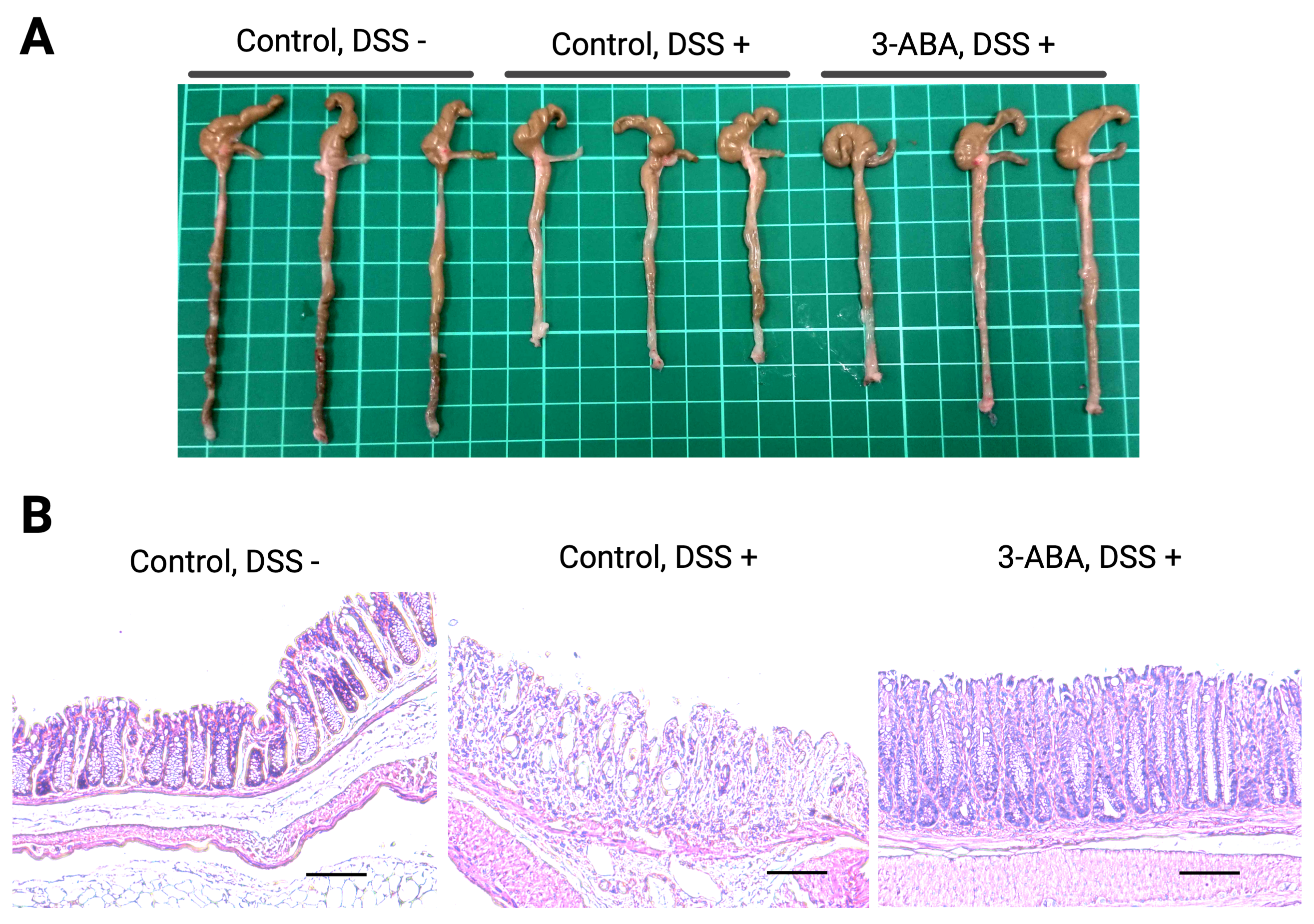

**Supplementary Figure 5.** Macroscopic and microscopic inflammation in mice with DSS-induced colitis. A representative macroscopic photograph of the colon (A) and histopathological images of the rectum stained with Hematoxylin and Eosin (B) from DSS-induced colitis mice. Scale bar, 100 μm.

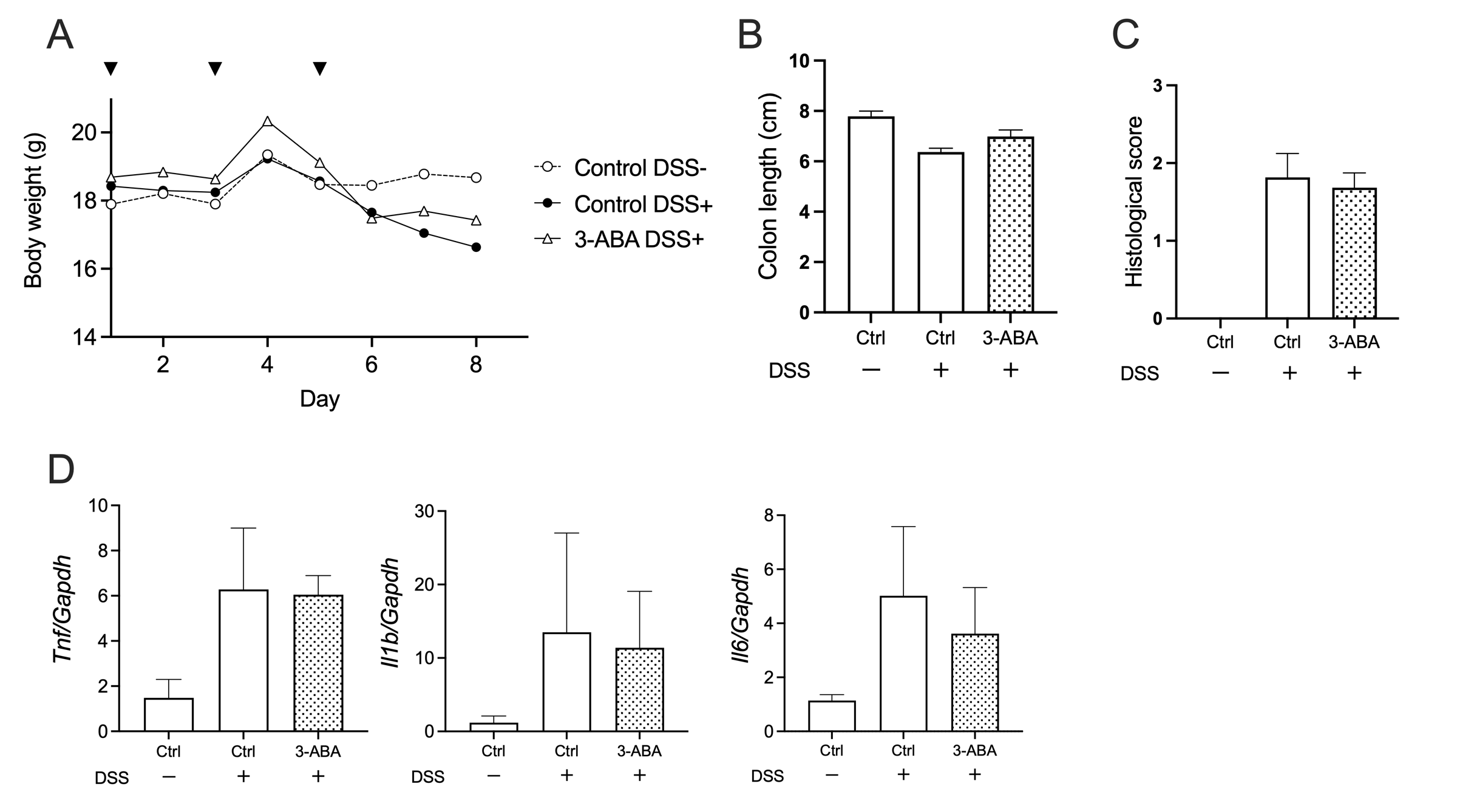

**Supplementary Figure 6.** Oral administration of 3-aminobenzoic acid attenuated DSS-induced colitis in mice. (A) Body weight change during the experiment. Arrowheads indicate the timing of oral drug administration. ^†^P<0.05 compared to the DSS-treated control mice (Control DSS+). The colon length (B), histologic score
(C), and gene expression of inflammatory cytokines (D) were evaluated on day 8. *P<0.05 **P<0.01. N=6 per group.

**Supplementary Table 1.** Clinical characteristics of patients with ulcerative colitis and healthy individuals.

| Disease | Age | Gender | Disease phenotype | Treatment | Endoscopic activity | Surgical History for UC |
| --- | --- | --- | --- | --- | --- | --- |
| NIBD | 36 | Male | - | - | - | - |
| NIBD | 40 | Male | - | - | - | - |
| NIBD | 37 | Male | - | - | - | - |
| NIBD | 45 | Male | - | - | - | - |
| NIBD | 34 | Male | - | - | - | - |
| NIBD | 35 | Female | - | - | - | - |
| UC | 34 | Female | Extensive colitis | Mesalazine | MES 3 | No |
| UC | 54 | Male | Extensive colitis | Mesalazine | MES 3 | No |
| UC | 32 | Female | Extensive colitis | Mesalazine | MES 2 | No |
| UC | 19 | Female | Extensive colitis | Mesalazine | MES 2 | No |
| UC | 30 | Male | Extensive colitis | Infliximab, Measalazine | MES 3 | No |

UC, ulcerative colitis; NIBD, healthy non-inflammatory bowel individuals; MES, Mayo endoscopic sub-score

**Supplementary Table 2.** List of primers and sequences for RT-qPCR

| Gene symbol | Forward primer | Reverse primer |
| --- | --- | --- |
| TJP1 | CGCTCTCGGGAGATGTTTATTTG | TCCTCCATTGCTGTGCTCTTG |
| CLDN1 | CTGCTGCTTCTCTCTGCCTT | GCTGGAAGGTGCAGGTTTTG |
| GAPDH | GATTTGGTCGTATTGGGCGC | TTCCCGTTCTCAGCCTTGAC |
| mTnf | AGGGGATTATGGCTCAGGGT | GAGTCCTTGATGGTGGTGCA |
| mIl1b | GGGCTGCTTCCAAACCTTTG | AAGACACAGGTAGCTGCCAC |
| mIl6 | TGGGGATGTCTGTAGCTCATTC | GCAACTGGATGGAAGTCTCTTG |
| mGapdh | ACTTTGGCATTGTGGAAGGG | TGGATGCAGGGATGATGTTCTG |
| mabA | AGCTCATCGAAGTCTGTCAGG | GGCTTTCGAACTTCTGCTCATC |
| mabB | ACGCCTATCTGTTCTCCATCAC | ATAGGCTTGCTCACGGTAGATG |
| 16S rRNA V4 | GTGCCAGCMGCCGCGGTAA | GGACTACHVGGGTWTCTAAT |

**Supplementary Table 3.** The percentage changes of TEER (ΔTEER%) after stimulation with 119 gut microbial metabolites in Caco2 cell monolayers.

| **ID** | **ΔTEER (%)** | **Name** | **CAS** |
| --- | --- | --- | --- |
| 1 | -15.10 | D-(-)-Lactic acid | 10326-41-7 |
| 2 | -7.60 | p-Methylphenyl potassium sulfate | 91978-69-7 |
| 3 | 5.90 | 3-Hydroxyhippuric acid | 1637-75-8 |
| 4 | 15.03 | N-Acetylputrescine hydrochloride | 18233-70-0 |
| 5 | 13.44 | Homogentisic acid | 451-13-8 |
| 6 | -27.00 | 4-Pyridoxic acid | 82-82-6 |
| 7 | -24.08 | Sphingomyelin | 85187-10-6 |
| 8 | -18.12 | Trimethylamine N-oxide | 1184-78-7 |
| 9 | -2.97 | Glycodeoxycholic Acid | 360-65-6 |
| 10 | -3.66 | 2,3-Butanediol | 513-85-9 |
| 11 | -5.73 | L-Gulose | 6027-89-0 |
| 12 | 9.74 | Allyl methyl sulfide | 10152-76-8 |
| 13 | -21.90 | Dimethyl trisulfide | 3658-80-8 |
| 14 | 16.10 | Norepinephrine | 51-41-2 |
| 15 | -4.61 | Norepinephrine (hydrochloride) | 329-56-6 |
| 16 | 9.41 | Folic acid | 59-30-3 |
| 17 | 11.87 | 3-Indoleacetic acid | 87-51-4 |
| 18 | 11.37 | (S)-Leucic acid | 13748-90-8 |
| 19 | 0.61 | D-Alanine | 338-69-2 |
| 20 | -13.33 | 2-Hydroxyhexanoic acid | 6064-63-7 |
| 21 | 13.70 | Cinnamoylglycine | 16534-24-0 |
| 22 | 8.72 | Thiamine monochloride | 59-43-8 |
| 23 | -3.70 | Melatonin | 73-31-4 |
| 24 | 10.24 | Niacin | 59-67-6 |
| 25 | 31.05 | L-Ascorbic acid | 50-81-7 |
| 26 | -9.89 | L-Ascorbic acid (sodium salt) | 134-03-2 |
| 27 | 6.70 | Salicylic acid | 69-72-7 |
| 28 | -3.31 | Sodium Salicylate | 54-21-7 |
| 29 | -12.70 | Lithocholic acid | 434-13-9 |
| 30 | -7.65 | Vitamin B12 | 68-19-9 |
| 31 | -4.11 | Creatinine | 60-27-5 |
| 32 | -78.90 | Biotin | 58-85-5 |
| 33 | 20.56 | 4-Aminobenzoic acid | 150-13-0 |
| 34 | 16.81 | Spermidine | 124-20-9 |
| 35 | -8.43 | Spermidine (trihydrochloride) | 334-50-9 |
| 36 | -49.57 | Spermine | 71-44-3 |
| 37 | -4.65 | Taurodeoxycholic acid (sodium hydrate) | 110026-03-4 |
| 38 | 27.41 | Tryptamine | 61-54-1 |
| 39 | -76.12 | Thiamine nitrate | 532-43-4 |
| 40 | -23.37 | Tartaric acid (disodium dihydrate) | 6106-24-7 |
| 41 | 12.89 | γ-Aminobutyric acid | 56-12-2 |
| 42 | -2.60 | Phloretin | 60-82-2 |
| 43 | 14.48 | Hyodeoxycholic acid | 83-49-8 |
| 44 | -7.01 | Indole-3-butyric acid | 133-32-4 |
| 45 | 10.45 | L-Phenylalanine | 63-91-2 |
| 46 | -76.59 | Benzoic acid | 65-85-0 |
| 47 | -3.28 | Protocatechuic acid | 99-50-3 |
| 48 | 9.97 | Cholesterol | 57-88-5 |
| 49 | -1.58 | Syringic acid | 530-57-4 |
| 50 | 5.05 | D-Mannitol | 69-65-8 |
| 51 | 11.33 | Homovanillic acid | 306-08-1 |
| 52 | 1.85 | Succinic acid | 110-15-6 |
| 53 | -1.67 | L-Lysine | 56-87-1 |
| 54 | 40.37 | L-Lysine hydrochloride | 657-27-2 |
| 55 | -1.89 | Tyrosol | 501-94-0 |
| 56 | -77.95 | Gallic acid | 149-91-7 |
| 57 | 16.57 | Gallic acid (hydrate) | 5995-86-8 |
| 58 | -8.73 | Allantoin | 97-59-6 |
| 59 | 6.78 | Hydroxytyrosol | 10597-60-1 |
| 60 | 17.12 | Deoxycholic acid | 83-44-3 |
| 61 | -1.50 | Deoxycholic acid sodium salt | 302-95-4 |
| 62 | -14.92 | trans-Cinnamic acid | 140-10-3 |
| 63 | -8.35 | Thiamine (hydrochloride) | 67-03-8 |
| 64 | -30.38 | Pyridoxine (hydrochloride) | 58-56-0 |
| 65 | -13.35 | Vanillic acid | 121-34-6 |
| 66 | 11.90 | D-(+)-Trehalose | 99-20-7 |
| 67 | -40.87 | D-(+)-Trehalose dihydrate | 6138-23-4 |
| 68 | 28.26 | Rhamnose | 3615-41-6 |
| 69 | -10.27 | Rhamnose (monohydrate) | 10030-85-0 |
| 70 | -27.37 | Glycodeoxycholic acid (monohydrate) | 1079043-81-4 |
| 71 | -7.47 | Taurochenodeoxycholic acid (sodium salt) | 6009-98-9 |
| 72 | -74.81 | Pyrogallol | 87-66-1 |
| 73 | 5.86 | 4-Hydroxyphenylacetic acid | 156-38-7 |
| 74 | 1.71 | Glycochenodeoxycholic acid | 640-79-9 |
| 75 | -24.64 | Glycochenodeoxycholic acid (sodium salt) | 16564-43-5 |
| 76 | 9.83 | Dihydrocaffeic acid | 1078-61-1 |
| 77 | -6.60 | D-Arabitol | 488-82-4 |
| 78 | -1.59 | Phosphorylethanolamine | 1071-23-4 |
| 79 | -2.18 | (E)-m-Coumaric acid | 14755-02-3 |
| 80 | 12.70 | cis,cis-Muconic acid | 1119-72-8 |
| 81 | -27.02 | 3,4-Dihydroxybenzeneacetic acid | 102-32-9 |
| 82 | 1.00 | 3-Hydroxyphenylacetic acid | 621-37-4 |
| 83 | -8.57 | Indole | 120-72-9 |
| 84 | 3.95 | 2,5-Dihydroxybenzoic acid | 490-79-9 |
| 85 | 2.73 | 2,5-Furandicarboxylic acid | 3238-40-2 |
| 86 | 13.90 | 3-Hydroxybenzoic acid | 99-06-9 |
| 87 | 1.74 | Nonadecanoic acid | 646-30-0 |
| 88 | -78.32 | 5-Hydroxymethyl-2-furancarboxylic acid | 6338-41-6 |
| 89 | -20.30 | 3-(3-Hydroxyphenyl)propionic acid | 621-54-5 |
| 90 | -2.65 | 3-Methyl-2-oxobutanoic acid | 759-05-7 |
| 91 | -76.60 | Skatole | 83-34-1 |
| 92 | -11.45 | 5-Hydroxyindole-3-acetic acid | 54-16-0 |
| 93 | 1.71 | Glutaric acid | 110-94-1 |
| 94 | 41.99 | 2-(1H-Indol-3-yl)ethan-1-ol | 526-55-6 |
| 95 | -9.60 | Creatine | 57-00-1 |
| 96 | -51.80 | 2-Phenylethylamine | 64-04-0 |
| 97 | 3.07 | 2-Phenylethylamine (hydrochloride) | 156-28-5 |
| 98 | 23.18 | N-Acetyl-L-tryptophan | 1218-34-4 |
| 99 | 7.24 | 3-Methylbutanoic acid | 503-74-2 |
| 100 | -32.14 | 2-(2-Phenylacetamido)acetic acid | 500-98-1 |
| 101 | 6.97 | Methyl 2-(1H-indol-3-yl)acetate | 1912-33-0 |
| 102 | 13.49 | 3-Indolepropionic acid | 830-96-6 |
| 103 | -0.46 | Desaminotyrosine | 501-97-3 |
| 104 | 14.22 | Disodium succinate | 150-90-3 |
| 105 | -10.20 | (R)-3-Hydroxybutanoic acid (sodium) | 13613-65-5 |
| 106 | 2.75 | 5-Aminovaleric acid | 660-88-8 |
| 107 | -81.30 | Glycolic acid | 79-14-1 |
| 108 | 8.52 | Hippuric acid | 495-69-2 |
| 109 | 10.93 | DL-3-Phenyllactic acid | 828-01-3 |
| 110 | 12.63 | Phenylacetylglutamine | 28047-15-6 |
| 111 | 8.32 | (R)-3-Hydroxybutanoic acid | 625-72-9 |
| 112 | -2.10 | Oxalic Acid | 144-62-7 |
| 113 | 12.77 | 4-Hydroxybenzoic acid | 99-96-7 |
| 114 | -12.09 | Urea | 57-13-6 |
| 115 | 8.63 | L-Tartaric acid | 87-69-4 |
| 116 | -29.29 | N-Methylsarcosine | 1118-68-9 |
| 117 | -47.70 | Pyruvic acid | 127-17-3 |
| 118 | -0.87 | Hydrocinnamic acid | 501-52-0 |
| 119 | -6.05 | Dimethyl sulfone | 67-71-0 |
